# From Garden Soil to Bio-electricity: Utilizing Rhizosphere Diversity for Microbial Fuel Cell Operation

**DOI:** 10.64898/2026.05.07.723474

**Authors:** Jinmo Lee, Byung Hong Kim

## Abstract

This study investigates the potential of the garden rhizosphere as a source of electrochemically active bacteria (EAB) for operating microbial fuel cells (MFCs). We evaluated a diverse array of garden flora, including vegetables (lettuce, Chinese cabbage), flowering plants (August lily, peppermint), and woody species (pine, oak, ginkgo, and bush clover). Among the tested groups, MFCs inoculated with peppermint and ginkgo rhizosphere microbiotas exhibited the highest current densities within their respective categories, significantly outperforming control groups without plant components. 16S rRNA gene microbial community analysis revealed that the initial rhizosphere environment acts as a decisive selective pressure, shaping distinct anode biofilms based on plant types (herbaceous vs. woody). Despite these structural differences in microbial assembly, high current generation was achieved in both peppermint and ginkgo systems, suggesting a high degree of functional redundancy within the rhizosphere-derived consortia. These findings demonstrate that various garden ecosystems can serve as robust biological reservoirs for MFC operation, where diverse microbial configurations are capable of sustaining efficient bio-electrochemical energy conversion.

## 1. Introduction

A microbial fuel cell (MFC) is a bio-electrochemical system that harnesses the metabolic activity of electrochemically active bacteria (EAB) to convert chemical energy from electron donors directly into electrical energy. Since the pioneering report of current generation by the Fe(III)-reducing bacterium *Shewanella putrefaciens* (Kim et al., 1999), MFC technology has undergone extensive development (Dixit et al., 2025; Shanthi Sravan et al., 2021).

EAB are known to be ubiquitous in nature, thriving in diverse environments where anaerobic oxidation occurs. This versatility has led to the emergence of plant-MFCs (PMFCs), which integrate living plants into the electrochemical circuit (Chong et al., 2024; Freire et al., 2026; Maddalwar et al., 2023; Mohanakrishna et al., 2026; Rusyn et al., 2025). Most PMFCs are designed to utilize plant exudates—organic compounds secreted by roots—as the primary fuel (Greenman et al., 2024; Kuleshova et al., 2022), while some systems incorporate wastewater as a supplementary energy source (Kacmaz and Eczacioglu, 2024).

The rhizosphere, the narrow zone of soil influenced by root secretions, hosts a complex microbiota essential for the host plant’s health and stress resilience (Babalola et al., 2021; Bakker et al., 2013, 2018; Benaissa, 2024; Berg et al., 2017; Enebe and Babalola, 2018; Luo et al., 2026; Pantigoso et al., 2022; Zhalnina et al., 2018). Plants excrete a significant portion of their photosynthesized carbon as exudates to support this beneficial microbiota (Baetz and Martinoia, 2014; Chen and Liu, 2024; Dhungana et al., 2023; Hartman et al., 2009; Huang et al., 2014; Nivetha et al., 2026). The composition of these exudates—and consequently the microbial community structure—is highly dependent on plant species, soil types, and interactions with neighboring vegetation (de Sousa et al., 2022; Dey et al., 2012; Qu et al., 2024; Seitz et al., 2022; Stassen et al., 2021; Tian et al., 2020; Tomazelli et al., 2024; Ulbrich et al., 2022).

The performance of PMFCs inherently depends on the synergy between the specific plant exudates and the resulting rhizosphere microbiota. In these systems, complex organic matters like polysaccharides and proteins are depolymerized and fermented into simple metabolites, such as acetate, which are then utilized by EAB like *Geobacter*—the most identified genus in PMFC anodes (Freire et al., 2026; Rusyn, 2021). Given this mechanism, we hypothesized that diverse flora found in a typical garden environment could provide a sufficient organic substrate and a robust microbial seed for sustainable MFC operation.

In this study, we evaluated the rhizosphere microbiotas of various garden plants— ranging from vegetables and flowering herbs to woody trees—as inoculums to enrich EAB. To minimize environmental variables such as soil type, all samples were collected from a localized garden area (within a 50-meter radius). By focusing on the inherent electrochemical potential of common garden species, this research aims to simplify the selection of plant-microbe systems for large-scale bio-energy applications and provides a comparative analysis of the microbial communities formed across different plant types.

## 2. Materials and Methods

### 2.1 Collection and Preparation of Rhizosphere Inocula

Rhizosphere samples, including plant hairy roots and their associated soil, were collected within a 50-meter radius of the following coordinates: 37°35’57.8”N, 127°03’18.0”E.. This localized sampling strategy was implemented to minimize potential experimental biases arising from variations in soil types and other environmental conditions. The selected plant species represented diverse garden flora, categorized into vegetables, flowering plants, and woody species, to evaluate their respective capacities for sustaining electrochemically active bacteria. The inoculum was prepared suspending about 5 grams of rhizosphere sample into 20ml of the phosphate-buffered acetate medium.

### 2.2 Medium Composition and Preparation

A phosphate-buffered acetate medium was utilized as the standardized electrolyte and substrate for all experiments. This medium was prepared by dissolving 0.9 g of NaCl, 0.2 g of MgSO_4_·2H_2_O, 0.2 g of CaCl·2H_2_O, 1.0 g of NH_4_Cl, 10.0 g of yeast extract and 25.0 g of sodium acetate in 1.0 L of 0.1 M phosphate buffer. For the purpose of enrichment and periodic replenishment, a ten-fold concentrated stock medium was separately prepared by dissolving the same quantities of chemical ingredients in 100 mL of the same phosphate buffer.

### 2.3 Configuration of the MFC Reactor

The experiments were conducted using dual-chamber MFC reactors as previously described by Kim et al. (2019). Each reactor chamber possessed an approximate working volume of 50 mL (5 × 5 × 2cm). Graphite felt (Nara Cell-Tech Co., Seoul, Korea) served as the primary material for both the anode and the cathode. Titanium wires were pierced through the graphite electrodes to function as current collectors. In some experiments, a platinum-coated carbon paper (Gas Diffusion Electrode, 0.3mg.cm-2 40% Pt on Vulcan, Nara Cell-Tech Co.) was also employed as the cathode. For these platinum-based cathodes, the titanium wire current collector was securely attached using a silver-based conductive glue. A Nafion 117 membrane was positioned between the chambers to serve as the proton exchange separator.

### 2.4 MFC Operation and Enrichment Procedures

All assembled MFC reactors were initially tested for potential leakage by filling the chambers with 0.1 M phosphate buffer (pH 7.0) before the commencement of the experiments. To initiate the enrichment process, 5 mL of the rhizosphere sample suspension and 5 mL of the 10 times concentrated medium were added to the anode chamber, which was then filled to the final volume with the phosphate buffer. When graphite felt was used as the cathode, 5 mL of the cathode buffer was replaced with 1.0 M potassium ferricyanide solution to act as the electron acceptor (Wei et al., 2012). The cathode compartments were continuously aerated at a flow rate of approximately 50 ml min-1 using a commercial fish tank air pump. The reactors were initially operated under an open-circuit configuration, and a 100 ohm external resistor was connected between the anode and cathode once the rate of increase in open-circuit potential began to stabilize. Throughout the enrichment phase, 5 mL of the anode electrolyte was periodically replaced with an equivalent volume of 10 times concentrated medium to ensure a continuous supply of substrates.

### 2.5 Data Acquisition and Statistical Analysis

The electrical signals generated by the MFC reactors were monitored and recorded on a personal computer using a Keithley DAQ6510 Data Acquisition/Multimeter System (Tektronix, Inc., Beaverton, OR, USA). All experimental groups were operated in triplicate to ensure reproducibility, and the results were expressed as mean values accompanied by their respective standard deviations. To maintain the reliability of the data, any individual reactor exhibiting a performance deviation of more than 25% from the other two reactors in the same group was excluded from the final statistical calculations. This exclusion was based on the assumption that such significant discrepancies likely resulted from mechanical system failures or localized leaks within that specific reactor unit.

### 2.6 16S rRNA-based microbial community analysis of MFC anode biofilms

Anode biofilm samples were collected from graphite felt electrodes of microbial fuel cells (MFCs) inoculated with different rhizosphere sources. These samples were sent to a company (LAS, Gimpo, Republic of Korea) for the microbiome analyses according to the following procedures. Each electrode was sectioned (1 × 1 cm), and DNA was extracted using the DNeasy PowerSoil Pro Kit (QIAGEN, Hilden, Germany) according to the manufacturer’s instructions. DNA concentration and purity were evaluated using a Quantus Fluorometer (Promega, Madison, WI, USA) and a NanoDrop 2000 spectrophotometer (Thermo Fisher Scientific, Waltham, MA, USA). The V4 region of the 16S rRNA gene was amplified using universal primers (515F/806R), and sequencing was performed by LAS (Gimpo, Republic of Korea) on the Illumina MiSeq platform using a paired-end (300 bp) approach.

Raw sequencing data were processed using QIIME 2 (v2022.2). Quality filtering, denoising, chimera removal, and amplicon sequence variant (ASV) generation were conducted using the DADA2 algorithm. Taxonomic assignment was performed against the SILVA database (v138). To ensure analytical precision, ASVs classified as chloroplast or mitochondrial sequences were removed prior to downstream analyses. Relative abundance profiles were generated to characterize microbial community composition. Beta diversity was assessed using Bray–Curtis dissimilarity and visualized by principal coordinate analysis (PCoA), and statistical significance was evaluated using PERMANOVA. All data visualization and statistical analyses were conducted using MicrobiomeAnalyst in conjunction with R (v4.5.0) and RStudio.

## 3. Results and Discussion

### 3.1. Enrichment and Electrochemical Characterization of EAB from Herbaceous Rhizospheres

The initial phase of this study focused on the enrichment of electrochemically active bacteria (EAB) from the rhizosphere microbiotas of garden grasses and vegetables. To evaluate the enrichment efficiency, dual-chamber MFC reactors were inoculated with rhizosphere samples, while reactors containing only buffer (blank) and barren soil (control) were operated under identical conditions for comparison.

All MFCs inoculated with rhizosphere microbiotas exhibited a rapid increase in open-circuit potential (OCP), reaching a maximum range of 750–800 mV within 24 hours (Table 1). This initial rise in OCP under non-loading conditions reflects the rapid establishment of a potential difference across the electrodes, representing the “charging” phase driven by the early metabolic activity of the inoculated microbes. Once the OCP reached a stable plateau— indicating a dynamic equilibrium between potential development and loss—the reactors were transitioned to a closed-circuit mode by connecting a 100 Ω external resistor.

**Table 1.** Performance of MFCs and enrichment of EAB inoculated with herbaceous and woody plant rhizosphere microbiotas.

| Plant type | Inoculum | Cathode type | Maximum open circuit potential |  | Maximum close circuit current |  |
| --- | --- | --- | --- | --- | --- | --- |
|  |  |  | Time±SD (Hr) | Mean±SD (mV) | Time±SD (Hr) | Mean±SD (μA) |
| Garden<br>grasses | Blank* | Plain graphite<br>with<br>potassium<br>ferricyanide | 13.7±0.7 | 725.9±11.5 | 130.7±9.3 | 45.43±7.70 |
|  | Control** |  | 13.7±0.7 | 715.5±10.9 | 125.7±0.4 | 53.16±2.70<sup>#</sup> |
|  | August lily |  | 24.0±2.0 | 805.5±14.9 | 126.3±0.7 | 55.23±9.07 |
|  | Lettuce |  | 22.6±1.4 | 763.7±30.2 | 120.0±8.0 | 65.55±2.51<sup>#</sup> |
|  | Blank | Pt-coated<br>graphite | 35.8±0.3 | 626.8±67.7 | 182.6±37.5 | 45.1±2.30 |
|  | Lettuce |  | 36.1±0.4 | 596.4±40.4 | 168.9±16.0 | 52.5±9.4 |
|  | Chinese cabbage |  | 34.1±3.1 | 691.8±38.9 | 157.6±40.7 | 48.2±1.8<sup>#</sup> |
|  | Pepper mint |  | 32.9±1.9 | 643.2±14.0 | 194.2±0.3 | 82.1±1.4<sup>#</sup> |
| Woody<br>plants | Pine | Pt-coated<br>graphite | 13.6±6.0 | 377.1±52.6 | 145.7±8.6 | 56.20±4.67<sup>#</sup> |
|  | Oak |  | 16.6±0.8 | 388.1±30.2 | 117.7±8.9 | 59.37±7.72 |
|  | Ginko |  | 13.0±13.0 | 420.1±13.8 | 123.7±22.7 | 73.65±0.08<sup>#</sup> |
|  | Bush clover |  | 40.7±16.0 | 358.6±6.4 | 98.1±5.8 | 64.12±4.77 |

\*Buffer solution, \*\* Soil from barren ground. All experiments were performed in triplicate
<sup>#</sup>To maintain the reliability of the data, any individual reactor exhibiting a performance deviation of more than 25% from the other two reactors in the same group was excluded from the final statistical calculations.

Upon closing the circuit, a substantial drop in potential was observed as expected, followed by a gradual and steady recovery in current generation. This subsequent increase in current, which continued for approximately 130 hours, is attributed to the proliferation and stabilization of the EAB population on the anode surface. To ensure that the observed current was primarily limited by the oxidative activity at the anode rather than cathodic constraints, potassium ferricyanide was employed as a potent electron acceptor at the graphite felt cathode (Wei et al., 2012). The periodic replacement of the cathode oxidant resulted in transient increases in current, further confirming that the system was successfully monitoring the enrichment of the anodic biofilm.

The comparative performance data, summarized in Table 1, demonstrate that while the blank and barren soil control groups eventually developed a degree of electrochemical activity, their OCP development was significantly slower and their stable current values were lower than those of the rhizosphere-inoculated groups. This finding suggests that while EAB may be ubiquitously present in the general environment, the rhizosphere of garden plants provides a much denser and more active community of these microbes. Among the tested species, the MFCs inoculated with lettuce rhizosphere microbiota exhibited a higher stable current compared to those inoculated with August lily, with the difference exceeding the calculated standard deviations.

In accordance with our data integrity protocol, certain reactors that showed deviations greater than 25% from their group means were excluded from the final statistical analysis. Such outliers were treated as potential system failures to ensure that the reported values accurately represent the characteristic performance of each plant-derived inoculum.

### 3.2. Evaluation of Pt-Coated Carbon Paper for Enhanced Stability

In the previous experiment, it was demonstrated that rhizosphere microbiotas from lettuce and August lily could rapidly enrich EABs and generate higher currents compared to the control groups. However, the use of potassium ferricyanide as a cathode oxidant resulted in periodic current fluctuations during the solution replacement process. To achieve more stable and continuous operation, the second experiment employed platinum (Pt)-coated carbon paper as cathodes, facilitating a direct oxygen reduction reaction (ORR) without the need for chemical oxidant replacement.

As summarized in Table 1, the maximum open-circuit potentials (OCP) observed in this configuration were generally lower than those recorded in the ferricyanide-based system. This decrease in OCP can be attributed to the lower redox potential of the Pt-catalyzed oxygen reduction under the given experimental conditions compared to the high electrochemical potential of the 0.1 M ferricyanide/graphite felt system. Despite the lower absolute potential, all MFC reactors inoculated with garden rhizosphere samples consistently outperformed the blank reactors, confirming the robust electrogenic activity of the garden-derived microbes.

The current generation profiles varied significantly depending on the plant species used for inoculation. Interestingly, the current densities for the lettuce-inoculated MFCs were lower in this Pt-coated carbon paper-cathode setup than in the previous ferricyanide-based experiment, likely due to the different cathodic limitations. Among the newly tested species, peppermint-inoculated MFCs exhibited the most superior performance, generating significantly higher current than both the lettuce and Chinese cabbage groups. While the MFCs inoculated with Chinese cabbage produced slightly higher current than the blank reactors, their output remained lower than those of the lettuce and peppermint groups. These differences between species were statistically significant, as they consistently exceeded the calculated standard deviations.

Consistent with the data quality control measures established in the first experiment, outlier reactors, which showed extreme deviations from their respective group means, were excluded from the final statistical calculations to ensure the reliability of the comparative analysis.

### 3.3. Comparative Analysis of MFC Performance Inoculated with Woody Plant Rhizospheres

The next stage of this study evaluated the electrochemical potential of rhizosphere microbiotas from various woody plants, contrasting their performance with the herbaceous species analyzed in the previous experiments. To ensure environmental consistency, samples were collected from a hillside and garden area within 50 meters of the initial sampling sites. The woody species included pine (*Pinus densiflora*), oak (*Quercus acutissima*), and bush clover *(Lespedeza bicolor*) from independent locations, as well as ginkgo (Ginkgo biloba) from the garden environment. These experiments were conducted using the platinum-coated carbon paper to maintain operational stability.

As summarized in Table 1, the MFC reactors inoculated with woody plant rhizospheres exhibited distinct electrochemical profiles compared to those of the herbaceous plants. Notably, the time required to establish a stable OCP was generally longer, and the maximum OCP values were significantly lower than those observed in the first and second experiments. These findings suggest that the microbial communities associated with woody roots may have different initial populations of EAB or that the nature of their root exudates provides a less immediate substrate for rapid charging.

Among the woody species tested, significant variations in current generation were observed. MFCs inoculated with the rhizosphere microbiotas of ginkgo and bush clover generated higher stable currents compared to those inoculated with pine and oak. The relatively lower performance of pine and oak-derived inocula may be attributed to the specific chemical composition of their rhizospheric environments, which might harbor lower densities of EABs or contain inhibitory compounds. However, all woody plant groups still demonstrated higher electrochemical activity than the blank control reactors, confirming that woody rhizospheres also possess the fundamental capacity to serve as a biological seed for MFC operation, albeit with different kinetic characteristics than garden grasses and vegetables.

### 3.4. Microbial community structure associated with current generation in MFCs

Fig. 1 shows microbial community structure of MFC anode biofilms derived from different rhizosphere sources. Bray–Curtis dissimilarity-based beta diversity analysis revealed clear differences in microbial community structure among samples, with principal coordinate analysis (PCoA) showing that PC1 (45.3%) and PC2 (25.1%) together explained 70.4% of the total variance. The blank control formed a distinct cluster, indicating that rhizosphere inoculation plays a decisive role in shaping the anode biofilm community. Samples were further grouped according to plant type, with herbaceous (lettuce, peppermint) and woody (ginkgo, pine, bush clover) sources forming well-defined clusters along the PC1 axis. This separation was statistically supported by PERMANOVA (R^2^ = 0.651, p = 0.0167), indicating that microbial community structure is primarily governed by the ecological characteristics of the rhizosphere inoculum rather than by the level of current generation. Envfit analysis showed a relatively high explanatory power between current generation and community structure (R^2^ = 0.7615), although this relationship was not statistically significant (p = 0.1194). High-performing samples, including ginkgo and peppermint, were distributed in distinct regions of the ordination space, indicating that similar levels of current can be achieved by different microbial community configurations. These results indicate that plant-specific rhizosphere environments, including differences in root exudate composition, act as strong selective pressures shaping microbial community assembly (Haichar et. al., 2008).

**Fig. 1.**
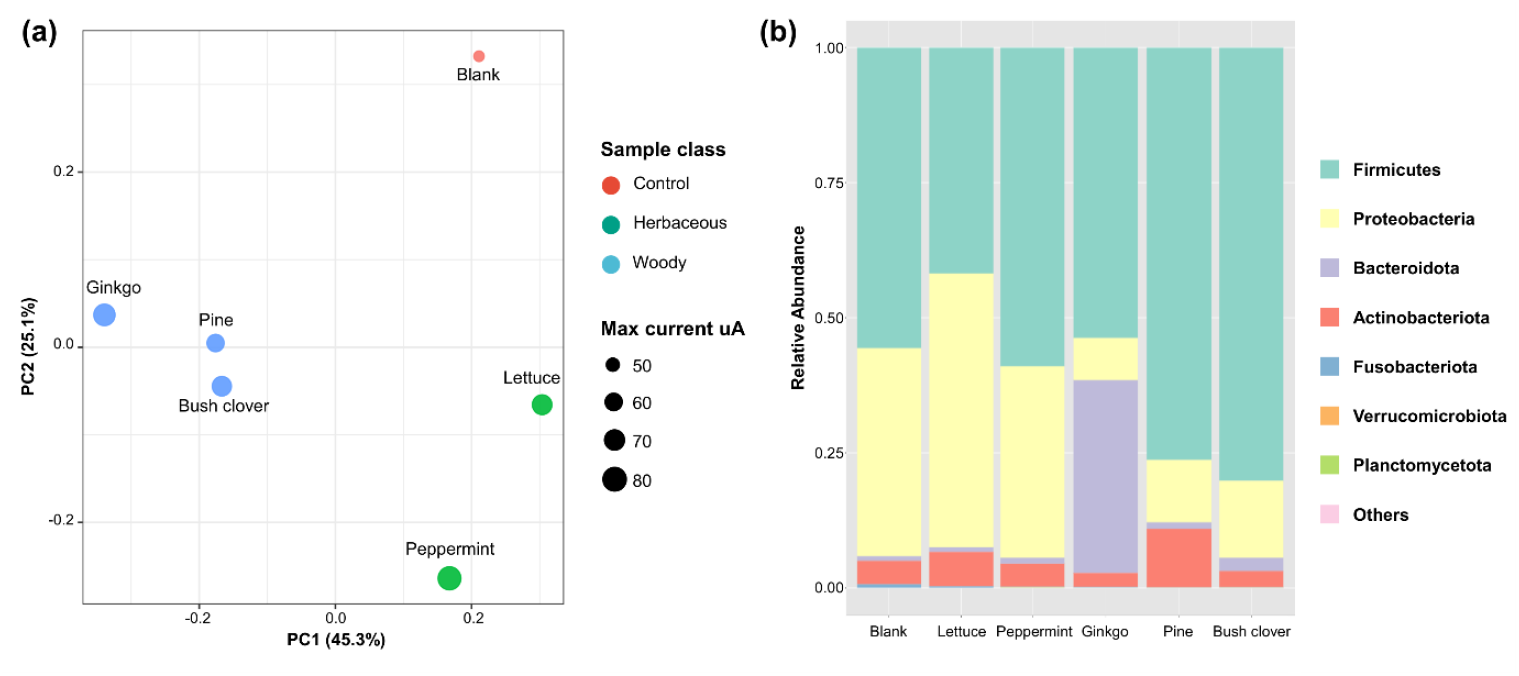
Microbial community structure of MFC anode biofilms derived from different rhizosphere sources. (a) Principal coordinate analysis (PCoA) based on Bray–Curtis dissimilarity showing clear separation of microbial communities according to plant type. (b) Relative abundance of dominant bacterial taxa at the phylum level across samples.

Relative abundance analysis at the phylum level supported these patterns. Firmicutes dominated across all samples, with Proteobacteria also representing a major component, consistent with microbial community structures commonly observed in MFC anode biofilms. Compositional differences were observed depending on the rhizosphere source. Lettuce samples exhibited a higher relative abundance of Proteobacteria, which includes representative electrochemically active bacteria such as *Geobacter*, suggesting the formation of a relatively balanced microbial consortium. Ginkgo samples showed an enrichment of Bacteroidota, while pine samples displayed a relative increase in Actinobacteriota, indicating selective enrichment of specific taxa. These findings indicate that, even under identical MFC operating conditions, the initial rhizosphere environment leads to the establishment of distinct microbial communities with different structural characteristics. The observation that high current generation occurred in systems with distinct community structures suggests that there is no single optimal microbial configuration for electricity production. Distinct microbial community structures appear capable of supporting electricity generation under different ecological configurations (Freguia et al., 2008; Yates et al., 2012; Jiang et al., 2018; Leicester et al., 2023).

## 4. Conclusions

The results of this study demonstrate that the rhizosphere microbiotas of various garden plants significantly enhance the performance of microbial fuel cells compared to barren soil and non-inoculated controls. The observation of low-level current generation in the control groups suggests that while EAB are ubiquitously distributed in the environment, the plant rhizosphere serves as a specialized niche that supports a substantially higher population and activity of these microbes. This confirms that the garden ecosystem can function as a natural reservoir of EAB, providing both the necessary microbial seeds and the organic substrates required for bio-electrochemical energy conversion.

Among the diverse flora tested, the rhizosphere microbiotas derived from peppermint (Mentha piperita) and the ginkgo tree (*Ginkgo biloba*) stood out for their superior current generation. This high electrochemical performance may be linked to the unique secondary metabolites and environmental adaptations of these species. Peppermint is well known for synthesizing menthol, a secondary metabolite with potent insect-repellent and antimicrobial properties. It is possible that such compounds selectively shape a rhizosphere microbiome that is enriched with robust, specialized bacteria capable of efficient extracellular electron transfer.

The performance of the ginkgo tree rhizosphere is particularly noteworthy. As one of the oldest living tree species, *Ginkgo biloba* is renowned for its extraordinary resilience to diseases, pests, and environmental stress. While the specific mechanisms behind its longevity and resistance are still being explored, our findings suggest that the ginkgo rhizosphere may host a highly efficient and stable community of electrogenic microbes. The ability of this “living fossil” to maintain a vigorous microbial ecosystem could be a key factor in its evolutionary endurance. The relatively higher current densities observed in ginkgo-inoculated MFCs compared to other woody species like pine and oak provide a new perspective on the biological robustness of this ancient tree.

Our microbiome analysis further substantiates these findings by revealing that the rhizosphere source acts as a decisive selective pressure in shaping the anode biofilm. Beta diversity analysis showed distinct clustering based on plant types—herbaceous versus woody—indicating that the initial ecological environment of the roots governs the microbial community assembly. Notably, the high current generation observed in both peppermint and ginkgo systems, despite their distinct microbial compositions, suggests a degree of functional redundancy where different microbial configurations can achieve similar levels of electrochemical output.

In conclusion, this research highlights the potential of using common garden flora as a sustainable source of microbial inocula for MFC technology. The distinctively high performance of peppermint and ginkgo rhizospheres, supported by their unique microbial community structures, underscores the versatility of garden ecosystems for bio-energy applications. By correlating these specific microbial configurations with electrochemical data, this study provides a foundation for the strategic selection of plant-microbe systems to optimize decentralized and sustainable energy production.

## Acknowledgement

Authors would like to express their deepest appreciations to Dr. J Shanthi Sravan (Gwangju Institute of Science and Technology) and Ha-Eun Cha (Sungkyunkwan University) for the help of MFC operation (JSS) and microbiome analysis (HEC) as well as training JL.

## Declaration of generative AI and AI-assisted technologies in the manuscript preparation process

During the preparation of this work, the authors used Gemini3.0 to improve the readability and language of the manuscript. After using this tool, the authors reviewed and edited the content as needed and take full responsibility for the content of the published article.

## References

Babalola, O. O., Emmanuel, O. C., Adeleke, B. S., Odelade, K. A., Nwachukwu Ayiti, B. C. O. E., Adegboyega, T. T., Igiehon, N. O., 2021. Rhizosphere microbiome cooperations: Strategies for sustainable crop production. Curr. Microbiol. 78, 1069–1085. 10.1007/s00284-021-02375-2

Baetz, U., Martinoia, E., 2014. Root exudates: the hidden part of plant defense. Trends Plant Sci. 19, 90–98. 10.1016/j.tplants.2013.11.006

Bakker, P., Berendsen, R. L., Doornbos, R. F., Wintermans, P. C. A., Pieterse, C. M. J., 2013. The rhizosphere revisited: root microbiomics. Front. Plant Sci. 4, 00165. 10.3389/fpls.2013.00165

Bakker, P. A. H. M., Pieterse, C. M. J., de Jonge, R., Berendsen, R. L., 2018. The soil-borne legacy. Cell 172, 1178–1180. 10.1016/j.cell.2018.02.024

Benaissa, A., 2024. Rhizosphere: Role of bacteria to manage plant diseases and sustainable agriculture - A review. J. Basic Microbiol. 64, 2300361. 10.1002/jobm.202300361

Berg, G. Köberl, M., Rybakova, D., Müller, H., Grosch, R., Smalla, K., 2017. Plant microbial diversity is suggested as the key to future biocontrol and health trends. FEMS Microbiol. Ecol. 93, fix050. 10.1093/femsec/fix050

Chen, L., Liu, Y. P., 2024. The function of root exudates in the root colonization by beneficial soil rhizobacteria. Biol.-Basel 13, 13020095. 10.3390/biology13020095

Chong, P. L., Chuah, J. H., Chow, C. O., Ng, P. K., 2024. Plant microbial fuel cells: A comprehensive review of influential factors, innovative configurations, diverse applications, persistent challenges, and promising prospects. Intern. J. Green Energy 22, 599–648. 10.1080/15435075.2024.2421325

de Sousa, L. P., Guerreiro, O., Mondego, J. M. C., 2022. The rhizosphere microbiomes of five species of coffee trees. Microbiol. Spect. 10 e00444-22. 10.1128/spectrum.00444-22

Dey, R., Pal, K. K., Tilak, K., 2012. Influence of soil and plant types on diversity of rhizobacteria. Proc. Nat. Acad. Sci., India Section B-Biological Sciences 82, 341–352. 10.1007/s40011-012-0030-4

Dixit, S., Imam, A., Rajput, R. S., Rajput, M. S., 2025. Microbial fuel cells as future of green technology: a systematic review. 3 Biotech 15, 218. 10.1007/s13205-025-04388-1

Dhungana, I., Kantar, M. B., Nguyen, N. H., 2023. Root exudate composition from different plant species influences the growth of rhizosphere bacteria. Rhizosphere 25, 100645. 10.1016/j.rhisph.2022.100645

Enebe, M. C., Babalola, O. O., 2018. The influence of plant growth-promoting rhizobacteria in plant tolerance to abiotic stress: a survival strategy. Appl. Microbiol. Biotechnol. 102, 7821–7835. 10.1007/s00253-018-9214-z

Freguia, S., Rabaey, K., Yuan, Z., Keller, J., 2008. Syntrophic processes drive the conversion of glucose in microbial fuel cell anodes. Environ. Sci. Technol. 42(21), 7937–7943. 10.1021/es800482e

Freire, J. B., Faustino, L., Abreu, A. A., Cruz, I. A. G. D. e., Costa, E. J. X., 2026. Microbial communities powering plant-microbial fuel cells: Diversity, functions and biotechnological perspectives. Microbial Biotechnol. 19, e70310, 10.scham.1111/1751-7915.70310

Greenman, J., Thorn, R., Willey, N., Ieropoulos, I., 2024 Energy harvesting from plants using hybrid microbial fuel cells; potential applications and future exploitation. Front. Bioeng. Biotechnol. 12, 76176. https://www.frontiersin.org/articles/10.3389/fbioe.2024.1276176

Haichar, F. E. Z., Marol, C., Berge, O., Rangel-Castro, J. I., Prosser, J. I., Balesdent, J., Achouak, W., 2008. Plant host habitat and root exudates shape soil bacterial community structure. ISME J. 2(12), 1221–1230. 10.1038/ismej.2008.80

Hartmann, A., Schmid, M., van Tuinen, D., Berg, G., 2009. Plant-driven selection of microbes. Plant and Soil 321(1-2), 235–257. 10.1007/s11104-008-9814-y

Huang, X. F., Chaparro, J. M., Reardon, K. F., Zhang, R. F., Shen, Q. R., Vivanco, J. M., 2014. Rhizosphere interactions: Root exudates, microbes, and microbial communities. Botany 92(4), 9. 10.1139/cjb-2013-0225

Jiang, Q., Xing, D., Zhang, L., Sun, R., Zhang, J., Zhong, Y., Ren, N., 2018. Interaction of bacteria and archaea in a microbial fuel cell with ITO anode. RSC Adv. 8(50), 28487–28495. 10.1039/C8RA01207E

Kacmaz, G. K., Eczacioglu, N., 2024. The mechanism of bioelectricity generation from organic wastes: soil/plant microbial fuel cells. Environ. Technol. Rev. 13(1), 76–95. 10.1080/21622515.2023.2283814

Kim, B. H., Kim, H. J., Hyun, M. S., Park, D. H., 1999. Direct electrode reaction of Fe (III)-reducing bacterium, Shewanella putrefaciens. J. Microbiol. Biotechnol. 9, 127–131. https://www.jmb.or.kr/journal/view.html?uid=554&vmd=Full

Kim, T., Yeo, J., Yang, Y., Kang, S., Paek, Y., Kwon, J. K., Jang, J. K., 2019. Boosting voltage without electrochemical degradation using energy-harvesting circuits and power management system-coupled multiple microbial fuel cells. J. Power Sourc. 410/411, 171–178. 10.1016/j.jpowsour.2018.11.010

Kuleshova, T., Rao, A., Bhadra, S., Garlapati, V. K., Sharma, S., Kaushik, A., Goswami, P., Sreekirshnan, T. R., Sevda, S., 2022. Plant microbial fuel cells as an innovative, versatile agro-technology for green energy generation combined with wastewater treatment and food production. Biomass and Bioenergy 167, 106629. 10.1016/j.biombioe.2022.106629

Leicester, D. D., Settle, S., McCann, C. M., Heidrich, E. S., 2023. Investigating variability in microbial fuel cells. Appl. Environ. Microbiol. 89(3), e02181–22. 10.1128/aem.02181-22

Luo, F., Cai, Y., Cui, Y., He, X., Xu, J., Tang, W., Wang, X., Cai, Y., Xie, H., Chen, W., Li, W., Ding, X., 2026. Microbiome eco-evolution of cultivated and wild rice species across the genus Oryza and its importance in supporting rice growth. BMC Microbiome 14, 83. 10.1186/s40168-026-02359-z

Maddalwar, S., Nayak, K. K., Singh, L., 2023. Evaluation of power generation in plant microbial fuel cell using vegetable plants. Bioresour. Technol. Reports 22, 101447. 10.1016/j.biteb.2023.101447

Mohanakrishna, G., Kudarimoti, V. I., Kamble, S. S., Naik, S. P., Manisha, S., 2026. Plant microbial fuel cells: A self-sustaining bioelectrochemical technology addressing sustainable development goals (SDGs) through bioelectricity production. Bioresour. Technol. 446, 134076. 10.1016/j.biortech.2026.134076

Nivetha, N., Asha, A. D., Kundu, A., Yadav, R. K., Abraham, G., Sangwan, S., Kumar, A., Paul, S., 2026. Mustard root exudates modulate early Bacillus spp. colonization under osmotic stress. J. Basic Microbiol. 66(1), e70115. 10.1002/jobm.70115

Pantigoso, H. A., Newberger, D., Vivanco, J. M., 2022. The rhizosphere microbiome: Plant– microbial interactions for resource acquisition. J. Appl. Microbiol. 133(5), 2864–2876. 10.1111/jam.15686

Qu, P., Wang, B. T., Qi, M. J., Lin, R., Chen, H. M., Xie, C., Zhang, Z. W., Qiu, J. C., Du, H. B., Ge, Y., 2024. Medicinal plant root exudate metabolites shape the rhizosphere microbiota. Intern. J. Mol. Sci. 25(14), 18. 10.3390/ijms25147786

Rusyn, I., 2021. Role of microbial community and plant species in performance of plant microbial fuel cells. Renew. Sustain. Energy Rev. 152, 111697. 10.1016/j.rser.2021.111697

Rusyn, I., Mittal, Y., Apollon, W., 2025. Plant microbial fuel cells: An innovative path toward integrated food and energy production for a sustainable future. J. Power Sour. 656, 238068. 10.1016/j.jpowsour.2025.238068

Seitz, V. A., McGivern, B. B., Daly, R. A., Chaparro, J. M., Borton, M. A., Sheflin, A. M., Kresovich, S., Shields, L., Schipanski, M. E., Wrighton, K. C., Prenni, J. E., Alexandre, G., 2022. Variation in root exudate composition influences soil microbiome membership and function. Appl. Environ. Microbiol. 88(11), e00226–22. https://journals.asm.org/doi/abs/10.1128/aem.00226-22

Shanthi Sravan, J., Tharak, A., Annie Modestra, J., Chang, I. S., Venkata Mohan, S., 2021. Emerging trends in microbial fuel cell diversification-Critical analysis. Bioresour. Technol. 326, 124676. 10.1016/j.biortech.2021.124676

Stassen, M. J. J., Hsu, S. H., Pieterse, C. M. J., Stringlis, I. A., 2021. Coumarin communication along the microbiome-root-shoot axis. Trends Plant Sci. 26(2), 169–183. 10.1016/j.tplants.2020.09.008

Tian, T., Reverdy, A., She, Q., Sun, B., Chai, Y., 2020. The role of rhizodeposits in shaping rhizomicrobiome. Environ. Microbiol. Rep. 2020 Vol. 12(2), 160–172. 10.1111/1758-2229.12816

Tomazelli, D., Peron, R. A. D., Mendes, S. D. C., Pinto, C. E., Baldissera, T. C., Baretta, D., Mendes, L. W., Goss-Souza, D., Klauberg, O., 2024. Plant diversity and root traits shape rhizosphere microbial communities in natural grasslands and cultivated pastures. Rhizosphere 29, 100864. 10.1016/j.rhisph.2024.100864

Ulbrich, T. C., Rivas-Ubach, A., Tiemann, L. K., Friesen, M. L., Evans, S. E., 2022. Plant root exudates and rhizosphere bacterial communities shift with neighbor context. Soil Biol. Biochem. 172, 108753. 10.1016/j.soilbio.2022.108753

Wei, L., Han, H., Shen, J., 2012. Effects of cathodic electron acceptors and potassium ferricyanide concentrations on the performance of microbial fuel cell. Intern. J. Hydrogen Energy 37(17), 12980–12986. 10.1016/j.ijhydene.2012.05.068

Yates, M. D., Kiely, P. D., Call, D. F., Rismani-Yazdi, H., Bibby, K., Peccia, J., Logan, B. E., 2012. Convergent development of anodic bacterial communities in microbial fuel cells. ISME J. 6(11), 2002–2013. 10.1038/ismej.2012.42

Zhalnina, K., Louie, K. B., Hao, Z., Mansoori, N., da Rocha, U. N., Shi, S., Cho, H., Karaoz, U., Loqué, D., Bowen, B. P., Firestone, M. K., Northen, T. R., Brodie, E. L., 2018. Dynamic root exudate chemistry and microbial substrate preferences drive patterns in rhizosphere microbial community assembly. Nature Microbiol. 3(4), 470–480. 10.1038/s41564-018-0129-3

